# Fitting side-chain NMR relaxation data using molecular simulations

**DOI:** 10.1101/2020.08.18.256024

**Authors:** Felix Kümmerer, Simone Orioli, David Harding-Larsen, Falk Hoffmann, Yulian Gavrilov, Kaare Teilum, Kresten Lindorff-Larsen

## Abstract

Proteins display a wealth of dynamical motions that can be probed using both experiments and simulations. We present an approach to integrate side chain NMR relaxation measurements with molecular dynamics simulations to study the structure and dynamics of these motions. The approach, which we term ABSURDer (Average Block Selection Using Relaxation Data with Entropy Restraints) can be used to find a set of trajectories that are in agreement with relaxation measurements. We apply the method to deuterium relaxation measurements in T4 lysozyme, and show how it can be used to integrate the accuracy of the NMR measurements with the molecular models of protein dynamics afforded by the simulations. We show how fitting of dynamic quantities leads to improved agreement with static properties, and highlight areas needed for further improvements of the approach.

## Introduction

Proteins are dynamical entities and a detailed understanding of their function and biophysical properties requires an accurate description of both their structure and dynamics. Nuclear magnetic resonance (NMR) experiments, in particular, have the ability to probe and quantify dynamical properties on a wide range of time scales and at atomic resolution. Computationally, molecular dynamics (MD) simulations also make it possible to probe both the structure and dynamics of proteins. Indeed, NMR and MD simulations may fruitfully be combined in a number of ways (***Case, 2002***; ***Lindorff-Larsen et al., 2005***), including using experiments for validating or improving the force fields used in simulations (***Norgaard et al., 2008***; ***Li and Brüschweiler, 2010**,**2011***), or for using the simulations as a tool to interpret the experiments (***Nodet et al., 2009***; ***Brookes and Head-Gordon, 2016***; ***Chen et al., 2019***; ***Vasile and Tiana, 2019***; ***Bottaro et al., 2020***).

A particularly tight integration between NMR and MD simulations has involved NMR spin relaxation experiments, which probe motions on the ps-to-ns timescales. Indeed, already soon after the first reported MD simulation of a protein (***McCammon et al., 1977***), such simulations were compared to NMR order parameters (***Lipari et al., 1982***). Integration between spin relaxation and MD is aided by the fact that MD can probe the relevant time scales and that the physical processes leading to spin relaxation are relatively well understood, and can therefore be modelled computationally.

The capability to reach given timescales does not, however, automatically translate into a perfect match between experiments and the simulated observables (***van Gunsteren et al., 2018***; ***Bottaro and Lindorff-Larsen, 2018***). Indeed, significant discrepancies between the two can emerge because of multiple factors, such as insufficient sampling (***Bernardi et al., 2015***), inaccurate description of the observable, i.e. the so-called forward model (***Cordeiro et al., 2017***), the imperfection of the underlying force fields (***Rauscher et al., 2015***; ***Henriques et al., 2015***; ***Robustelli et al., 2018***; ***Piana et al., 2020***; ***Nerenberg and Head-Gordon, 2018***) or a combination of these factors.

NMR spin relaxation experiments are generally interpreted using variants of the so-called model free formalism (***Halle and Wennerström, 1981***; ***Lipari and Szabo, 1982***; ***Clore et al., 1990***) that interprets the experimental data using generalized order parameters that describe the amplitudes of the motions and time scales associated with those motions. A common approach to compare MD and NMR relaxation experiments involves calculating order parameters from the simulations and comparing to those extracted from the experiments. The results of such comparisons have shown that while MD simulations generally give a relatively good agreement with order parameters for the backbone amide groups, the agreement for methyl bearing side chains is more variable (***Bremi et al., 1997***; ***Skrynnikov et al., 2002***; ***Best et al., 2005***; ***Showalter et al., 2007***; ***Liao et al., 2012***; ***O’Brien et al., 2016***; ***Bowman, 2016***; ***Anderson et al., 2020***).

A more detailed and richer combination, however, involves calculating the NMR relaxation rates directly from the simulations and comparing these to experiments. While this approach does not necessarily provide detailed information about the timescales of these motions, it has the advantage of not being dependent on a specific analytical model which might influence the interpretation of the relaxation data. Recently, an approach was developed to calculate deuterium NMR relaxation rates in methyl bearing side chains from MD simulations (***Hoffmann et al., 2018b, 2020***). Comparison to experiments revealed systematic deviations from simulations with several different force fields, and corrections to the methyl torsion potential were developed based on quantum calculations and shown to increase agreement with experiments (***Hoffmann et al., 2018a**, **2020***). Despite these improvements and a substantial body of work on studies of side chain dynamics (***Lee et al., 1997***; ***Palmer III, 1997***; ***Ming and Brüschweiler, 2004***; ***Cousin et al., 2018***; ***Anderson et al., 2020***), the agreement remains imperfect and further improvements would be desirable.

One possible way to reduce the systematic error coming from force field inaccuracies is to introduce a bias in the ensemble generated by MD simulations to improve agreement with the experimental data. Such a bias may either be introduced during the simulation by adding a correction to the force field that depends on the experimental measurements, or after the simulation has been completed through a procedure known as *reweighting*. Using biased simulations it is, for example, possible to construct conformational ensembles that are in agreement with backbone and side chain order parameters from NMR spin relaxation (***Best and Vendruscolo, 2004***; ***Lindorff-Larsen et al., 2005***).

Most implementations of such methods for biasing against experiments work by updating some ‘prior’ information, typically encoded in the MD force field, with information from the experiments by changing the weights of each conformation so as to improve the agreement with experiments when calculated with these new weights. Often these problems are highly underdetermined in that there are many more parameters (weights) to be determined than experimental measurements. Thus, an important ingredient in many methods is a framework to avoid overfitting by balancing the information from the force field with that from the data, and many approaches apply Bayesian or Maximum Entropy formalisms for this purpose (***Bonomi et al., 2017***; ***Orioli et al., 2020***).

Although different methods do so in different ways and with different assumptions, most reweighting methods share a key commonality: they can only deal with static (equilibrium) observables, i.e. quantities that can be expressed as ensemble averages over a set of configurations (***Bonomi et al., 2017***; ***Orioli et al., 2020***). Evidently, this complicates the generation of conformational ensembles that match the spin relaxation data, which depend both on the amplitudes and time scales of the motions.

A few approaches have been described to fit NMR relaxation data directly. For example, the isotropic reorientational eigenmode dynamics (iRED) approach (***Prompers and Brüschweiler, 2002***) uses a covariance matrix to calculate spin relaxation, and may be applied to blocked simulations thus indirectly taking into account the time scales of the motions (***Gu et al., 2014***). Recently, a more direct approach has been described to reweight MD simulations against NMR spin relaxation data (***Salvi et al., 2016**, **2019***). The basic idea is to split one or more simulations into smaller parts (blocks), each of which are long enough to contain information about both the amplitudes and time scales of the motions that lead to spin relaxation. By averaging over such blocks, one can then estimate NMR relaxation parameters that may be compared to experiments. Similar to the reweighting methods described above for individual configurations, one can instead use reweighting of the blocks and thus determine weights that improve agreement with the experimental data without having to decompose it into order parameters. This approach, called ABSURD (Average Block Selection Using Relaxation Data), has been applied to the analysis of backbone NMR relaxation of intrinsically disordered proteins that are difficult to analyse using conventional model free techniques.

We here describe a new approach to interpret NMR relaxation data, focusing our application on side chain dynamics. Our approach builds upon the ABSURD method, and includes two extensions. First, we use the recently described methods to calculate side chain NMR relaxation parameters from molecular dynamics simulations (***Hoffmann et al., 2018b***). Second, we extend ABSURD by including an entropy restraint term in the optimization that helps avoid overfitting. We thus term our method ABSURDer (Average Block Selection Using Relaxation Data with Entropy Restraints). We applied ABSURDer to study the fast timescale dynamics of T4 lysozyme (T4L), a protein which has been the subject of numerous experimental and computational studies. We validate and describe our method using synthetic data and then apply it to data from NMR experiments. We find that reweighting simulations with ABSURDer improves the agreement between methyl relaxation rates in synthetic data sets coming from both the same and a different force field. Furthermore, we find that these reweighted trajectories also show improvements in the agreement of equilibrium observables, such as rotamer distributions. Finally, we use experimental NMR data for reweighting to improve agreement between simulations and experiments. In this case we find smaller improvements are possible, and we discuss possible origins of this observation.

## Results and Discussion

### Overview of the ABSURDer approach

The workflow in ABSURDer consists of three separate steps (Fig. 1). Steps 1 and 2 correspond to a previously described method for calculating side chain relaxation, and step 3 corresponds to the ABSURD approach with the inclusion of an additional restraint to decrease the risk of overfitting, and to balance the information in the force field with that in the data.

**Figure 1.**
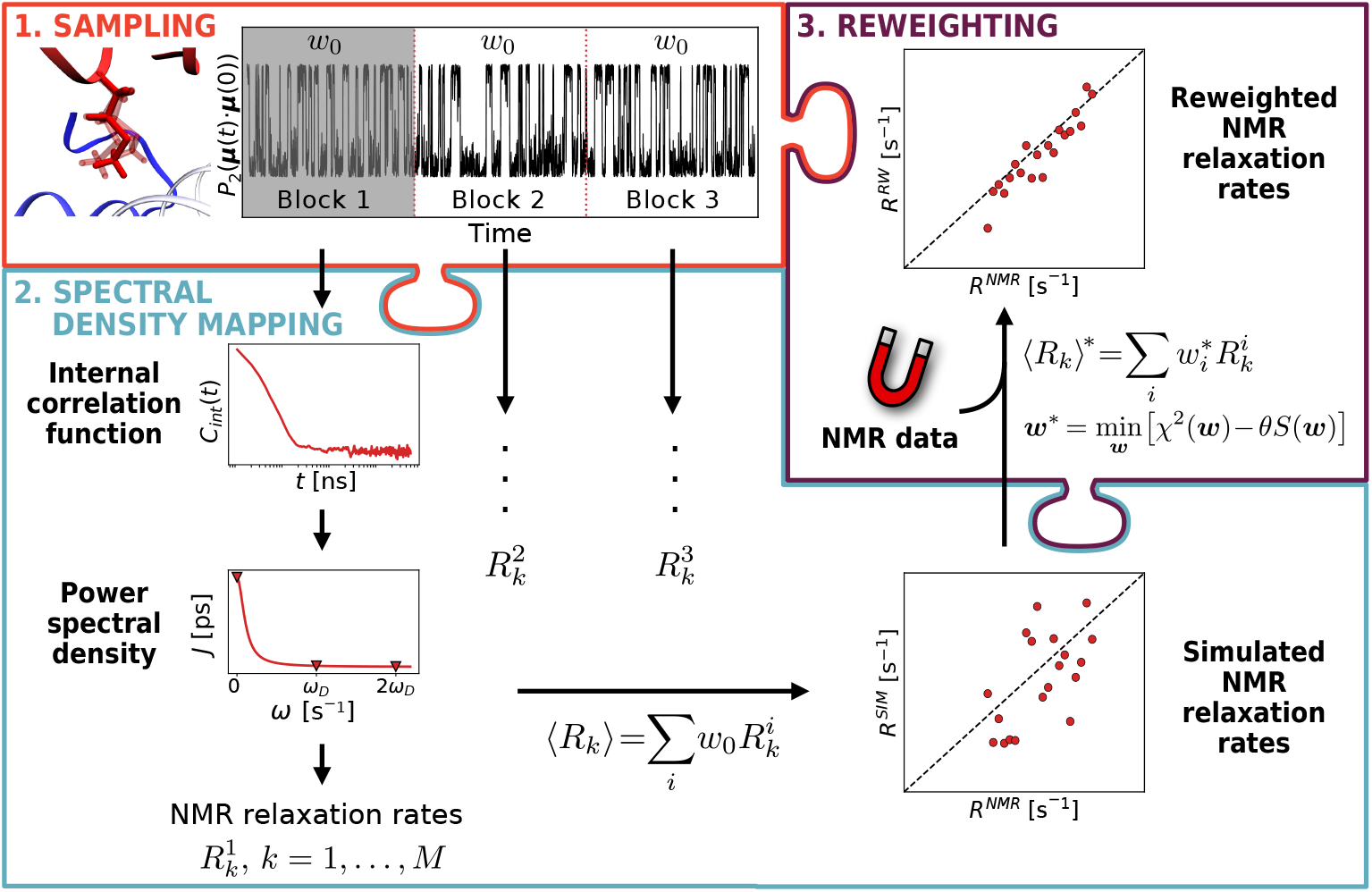
Schematic representation of the ABSURDer workflow. Steps 1 involves sampling protein dynamics using one or more longer MD simulations and calculating bond vector orientations for the resulting trajectories. These are then divided into blocks and in Step 2 we calculate correlation functions, spectral densities and NMR relaxation rates for each block. In Step 3, we optimize the agreement between the average calculated rates and experimentally measured values by changing the weights of the different blocks. For details see main text, methods and code online.

First, we performed three independent 5 μs-long MD simulations to sample the structure and dynamics of T4L. We here use the recently optimized Amber ff99SB*-ILDN force field with the TIP4P/2005 water model and a modification of the methyl torsion potential, based on CCSD(T) coupled cluster quantum chemical calculations of isolated dipeptides, that improves agreement with NMR relaxation data (***Hoffmann et al., 2018a***). See the Methods section for further details about all simulations.

Second, we divided the total of 15 μs simulations into 1500 10 ns blocks, and calculated NMR relaxation parameters for each of these independently (see Methods, and below for a more detailed discussion on the origin of this block size). Briefly described we: (i) calculate internal correlation functions ***C_int_***(***t***) for methyl C-H bonds; (ii) use backbone motions to describe the global overall tumbling motion; (iii) combine internal and global rotational motions to calculate the spectral density function ***J***(***ω***) and (iv) extract NMR relaxation parameters from this. We here focus on three NMR relaxation parameters ***R***(***D**_z_*), ***R***(***D**_y_*) and 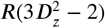, and calculate these rates for each methyl group and each of the 1500 blocks. As done previously (***Hoffmann et al., 2018b***) we exclude residue ALA146 from the analyses.

The third step in our approach consists of reweighting the MD simulations to improve agreement with the experimental data. Our approach combines the ABSURD method with concepts from Bayesian/Maximum Entropy ensemble refinement. The goal here is to assign to each of the 1500 10 ns simulations a different weight (*w_i_*; *i* = 1,…, 1500) so as to improve agreement with the experimental data, as quantified by the *χ^2^* between experimental and calculated NMR relaxation rates. Such an approach is, however, prone to overfitting and does not necessarily utilize the information about the conformational landscape encoded in the molecular force field. Thus, we and others have previously described methods that circumvent this problem by adding an entropy regularization to the *χ*^2^ minimization (***Gull and Daniell, 1978***; ***Cesari et al., 2018**; **Köfinger et al., 2019***; ***Bottaro et al., 2020***; ***Orioli et al., 2020***), by instead minimizing the functional 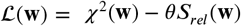 (see Methods). In this equation *S_rel_* represents the relative entropy that measures how different the weights are from the initial (uniform) weights, and thus how much the final set of simulations differs from the initial set. Typically, we represent this as *ϕ_eff_* = exp(*S_rel_*), which represents the effective fraction of the original 1500 blocks (sub-simulations) that contribute to the final ensemble. In these equations the parameter *θ* sets the balance between fitting the data (minimizing *χ*^2^) and keeping as much as possible of the original simulation (maximizing *ϕ_eff_*). We refer the reader to previous literature about the methods overall and how best to select this parameter (***Andrae et al., 2010***; ***Hummer and Köfinger, 2015***; ***Bottaro et al., 2018***; ***Cesari et al., 2018***; ***Köfinger et al., 2019***; ***Crehuet et al., 2019***; ***Chen et al., 2019***; ***Bottaro et al., 2020***; ***Orioli et al., 2020***).

To describe and understand how well ABSURDer works, we have applied it both to synthetic and experimental data for T4L. We used data of increasing complexity in an attempt to disentangle problems arising from the sampling, the force field, the forward model to calculate relaxation data and our approach to fit the data. In all cases, we employed the 1500 10 ns blocks coming from the three 5 μs simulations with the Amber ff99SB*-ILDN force field to fit the data. In our first application we created synthetic data using the same force field by running five additional 1 μs-long simulations. In this case, the only differences between the ‘data’ and the simulations would arise from insufficient sampling. In our second application, we again created synthetic data, in this case however using the Amber ff15ipq force field (with a modified methyl torsion potential (***Hoffmann et al., 2020***)) and the SPC/E_*b*_ water model (***Takemura and Kitao, 2012***; ***Debiec et al., 2016***). We performed three 1 μs simulations and used these to generate synthetic NMR relaxation data. This test enables us to examine how the method behaves where there are real differences between the potential used in the simulations we use to fit the data, and that give rise to the ‘experimental data’. In contrast to using actual experimental data, however, we here have access to the full underlying ensembles and dynamics, and thus are able to compare e.g. full spectral density function and rotamer distributions. Finally, we applied ABSURDer to experimental NMR relaxation data, providing an example of how the approach might be used in practice. The results of these three levels of complexity are described in the following sections.

### Determining block size and assessing convergence

The choice of the block length plays a central role in the calculation of the relaxation rates and in the possibility to optimize the results via reweighting. For this, two opposing effects need to be considered (Fig. 2A). On one hand, a long block size allows for a more correct representation of the correlation functions and thus estimation of the NMR relaxation rates. On the other hand, it is desirable to increase the variation across the different blocks, as this makes it easier to fit the relaxation data by reweighting; however this variance decreases with the length of the blocks (Fig. 2A). Ideally, one would choose a block length that is (i) long enough not to introduce too much bias compared to calculating correlation functions over the full trajectory and (ii) short enough that each block is not just a ‘converged’ representation of the full trajectory. We chose 10 ns as a block length that provides a large number of blocks to be used in the fit and still balances these two requirements (Fig. 2A). This choice allows us to calculate the internal time correlation functions of the methyl C-H bonds up to a maximum lag time of 5 ns, as also done previously (***Hoffmann et al., 2020***). We note that the chosen block size is also close to the global tumbling time of the protein (***τ_R_*** ≈ 11 ns). As NMR spin relaxation experiments are mostly sensitive to motions on time scales up to approximately ***τ_R_***, this explains why much shorter block sizes do not capture the NMR relaxation parameters accurately. Examining each of the three relaxation rates separately (Fig. S1) we find that in particular ***R***(***D**_y_*) is poorly determined using short blocks. Finally, we note that while calculations of correlation functions from just a single 10 ns of simulation is generally not sufficient to compare to NMR relaxation data, in all our analyses we average over large numbers of blocks. In that case, we find that averaging over 1500 10 ns blocks gives very similar results as 3 5 μs blocks (Fig. S2). This is indeed expected as the division into blocks will only affect the calculations of the correlation that ‘cross the boundary’ of the blocks.

**Figure 2.**
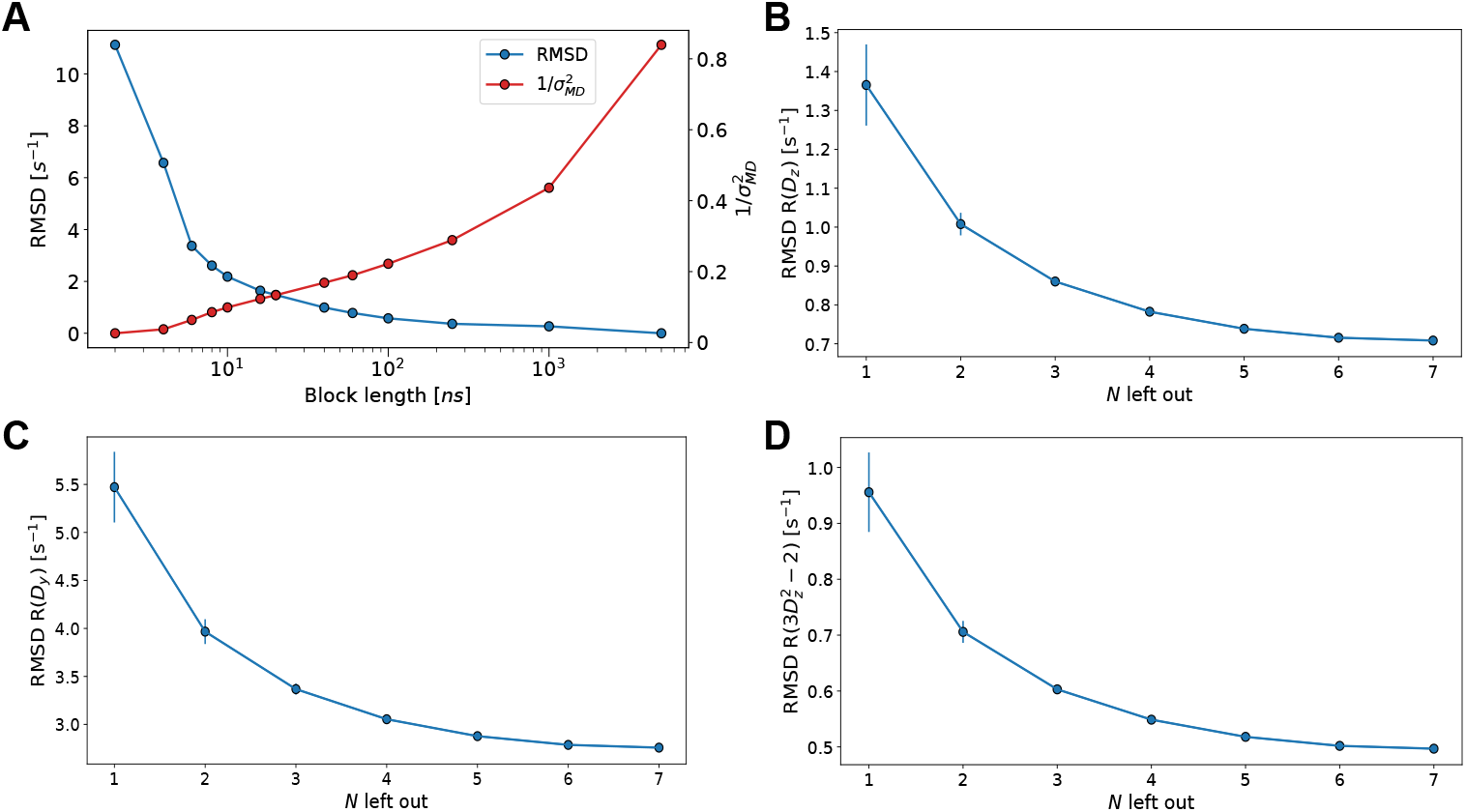
Choice of block size and convergence of NMR relaxation rates. (A) Overall (blue) RMSD of NMR relaxation rates and (red) average inverse variance with respect to the full MD data set as a function of the block length. (B-D) Average RMSD of the three NMR relaxation rates obtained by comparing a set of *N* 1 μs long segments to the rates calculated from the remaining 15 - *N* segments.

To assess the level of convergence, we divided our three 5 μs-long simulations into 15 1 μs segments and employed leave-*N*-out cross-validation to estimate the impact of the amount of sampling for the estimation of relaxation rates. For each value of *N*, we calculated the NMR relaxation rates by averaging over the *N* segments, and then compared to the average over the remaining 15 – *N* segments by calculating the root mean square deviation (RMSD) between the two sets of rates. We repeated these calculations for all possible combinations of leaving out *N* segments, and the RMSD values were averaged. Finally, we repeated these calculations for all values of *N* (*N* = 1,…, 7) (Fig. 2B-D). Even at *N* = 1 we find relative low RMSD values when compared to the spread of the rates (Fig. 2A-C), and these errors decrease further as additional sampling is included. Thus, we conclude that using several microseconds of sampling we are able to obtain relatively precise estimates of the relaxation rates, as would be expected given that they report on ps-ns dynamics in the protein.

### Fitting synthetic data generated with the same force field

We ran five 1 μs-long simulations with the Amber ff99SB*-ILDN force field and calculated NMR relaxation parameters from these. In what follows, this synthetic data will sometimes be referred to as ‘NMR’ or ‘experiments’ to indicate that we treat them as observables from an experiment. We calculated the same observables as the average over 1500 10 ns blocks coming from the three 5 μs simulations with the same force field, and compared them to the synthetic experimental data (Fig. 3A–C). As expected the calculated and experimental values are strongly correlated, in line with the fact that these were generated from the same underlying physical model. The calculated values of a reduced *χ*^2^ were 22, 15 and 17 for ***R***(***D**_z_*), ***R***(***D**_y_*) and 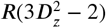, respectively.

**Figure 3.**
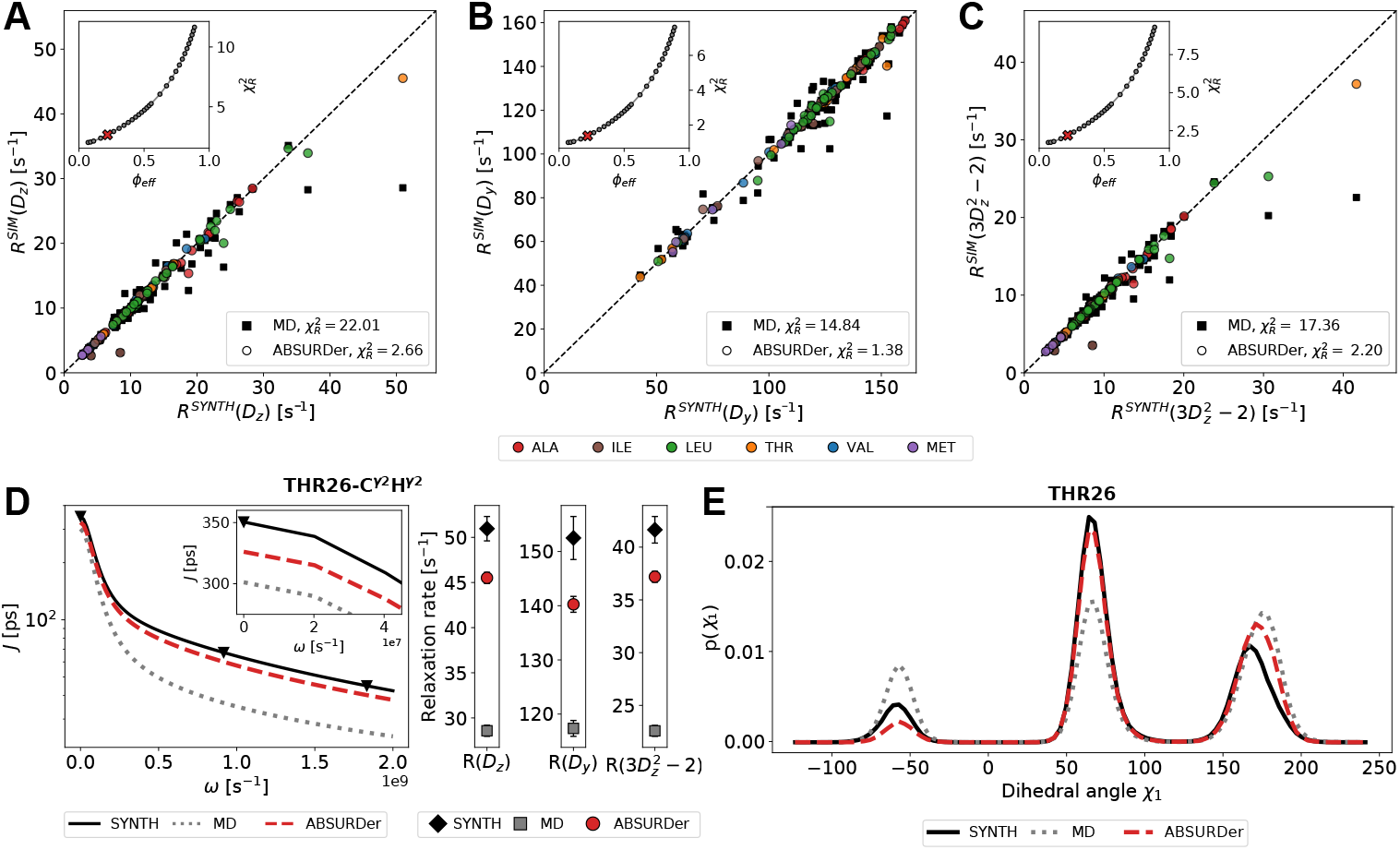
ABSURDer applied to synthetic data generated with the same force field. (A - C) Comparison between synthetic ‘experimental data’ and simulations for ***R***(***D**_z_*), ***R***(***D**_y_*) and 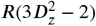 both before (black squares) and after (filled circles) reweighting with ABSURDer. The rates are coloured by residue type. The insets show the behaviour of the respective reduced *χ*^2^ vs. *ϕ_eff_* during the reweighting using different values of *θ*. The chosen *θ* = 300 (resulting in *ϕ*_eff_ ≈ 0.2) is shown as a red cross. (D)The spectral density function and ***R***(***D**_z_*), ***R***(***D**_y_*) and 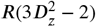 of THR26-C^*γ*2^H^*γ*2^ before (grey/dotted line) and after (red/dashed line) reweighting with ABSURDer. ***J***(0), ***J***(*ω*) and ***J***(2*ω*) are shown as black triangles. The errorbars represent the standard error of the mean from averaging over the 1500 10 ns sub-trajectories. (E) The *χ*_1_ angle probability density distribution of THR26 before (grey/dotted line) and after (red/dashed line) reweighting with ABSURDer.

We then asked whether we could improve the agreement between experiments and simulations by changing the weights of each of the 1500 blocks to reweight the calculated observables. We thus determined a set of weights that minimize the functional 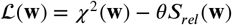 at different values of *θ* and plot the resulting *χ*^2^ vs. *ϕ_eff_* (Fig. 3A-C; insets). It is clear that as *θ* is decreased to put greater weight on fitting the data, the agreement between experiment and simulation can be improved substantially, though at the cost of utilizing only a fraction of the input simulations. For simplicity, we here opted to use a value *θ* that gives rise to *ϕ_eff_* ≈ 0.2, though our conclusions are relatively robust to this choice. At this level of fitting the agreement between experiment and simulations improves substantially (reduced *χ*^2^ approximately 3,1 and 2 for ***R***(***D**_z_*), ***R***(***D**_y_*) and 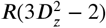, respectively). We note also that *ϕ_eff_* ≈ 0.2 represents a total of 3 μs of MD simulation, which in itself is sufficient to obtain relatively converged values (Fig. 2B-D).

In addition to reweighting the simulations using all three types of relaxation rates (***R***(***D**_z_*), ***R***(***D**_y_*) and 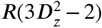), we also performed reweighting using each rate individually or in pairs, and used the remaining rate(s) for cross validation (Fig. S3). Overall, we find that fitting one or two rates leads to improvements also in rates that are not used in reweighting. For certain rates and very aggressive reweighting (low values of *ϕ*_eff_) we observe, however, an increase in the cross validated rates. This behaviour is expected as the three rates in part depend and report on the same properties of the spectral density function. Indeed, it is clear that ***R***(***D**_y_*) is less correlated with the two other rates, which is likely due to the fact that only this relaxation rate depends on the spectral density at zero frequency (***J***(0)). Nevertheless, the general improvement in cross validation down to *ϕ*_eff_ ≈ 0.2 provides another reason why we chose this value for our analyses.

While the NMR relaxation parameters are the experimental observables, they are calculated from the simulations via the spectral density functions, ***J***(*ω*). Spectral density functions are often estimated from experiments using simplified and analytically tractable functions that balance the complexity of the motions and the number of free parameters that can be estimated from limited experiments. Using synthetic data gives us the possibility of comparing the spectral density functions from the simulations, both before and after reweighting, with those that give rise to the experiments. We here exemplify this using a single methyl group from THR26 (Fig. 3D); the code and data to generate plots for all residues are available online (github.com/KULL-Centre/papers/tree/master/2020/ABSURDer-kummerer-orioli-et-al-2020). The observed NMR relaxation parameters depend on the spectral density at three frequencies (*ω* = 0, *ω_D_* and *ω*_2*D*_, respectively) and it is thus not surprising that fitting to this data improves agreement at these frequencies. Nevertheless, it is also clear that our reweighting of the trajectories leads to a general improvement between the calculated spectral density function and that which was used to generate the synthetic data. As it is clear from this example (Fig. 3D), and from the overall results on all rates (Fig. 3A-C), it is also evident that discrepancies between experiments and simulations remain. While it is possible to fit the data more closely by decreasing *θ* this comes with the risk of potential overfitting (Fig. S3) and diminished trust of the input simulations.

The observed NMR relaxation parameters depend on a complex combination of both overall tumbling and various types of internal motions. In addition to the fast rotation of the methyl groups, this includes fluctuations of the backbone and motions both within and between rotameric states. It has previously been shown that rotamer jumps, in particular when this occurs on a timescale faster than rotational tumbling, can contribute substantially to side chain relaxation (***Best et al., 2005***; ***Hu et al., 2005***). We thus calculated the side chain dihedral angles and compared the values before and after reweighting to those from the simulation used to generate the synthetic data. Highlighting again THR26 we see that, due to imperfect sampling of rotamer distributions, even in multi-microsecond simulations, there are clear differences between the distribution of the *χ*_1_ dihedral angle in the simulations used to generate the data compared to that used to fit it (Fig. 3E). Particularly important, however, we also see a clear improvement after the reweighting. Thus, we find that by reweigting a ‘dynamic’ property (NMR relaxation rates) we also find improvement in a ‘static’ property such as the distributions of dihedral angles. Similar results were found for a wide range of dihedral angles including also in the backbone (Fig. S5) (see also github.com/KULL-Centre/papers/tree/master/2020/ABSURDer-kummerer-orioli-et-al-2020).

### Fitting synthetic data generated with a different force field

The calculations above demonstrate how ABSURDer can be used to fit NMR data using MD simulations, but neglects the challenges arising from imperfect force fields. Thus, we increased the complexity of our comparisons by generating synthetic ‘experimental’ data with a different force field than that used to fit the data. We thus ran three 1 μs-long simulations with the Amber ff15ipq force field and calculated synthetic NMR relaxation parameters from these. We again used ABSURDer to fit the 1500 10 ns blocks to this data. In this case, imperfect agreement arises both due to insufficient sampling and differences in the underlying potential, leading to differences both in the dynamics and thermodynamic averages.

This added complexity is evident when we compare ‘experimental’ and calculated NMR relaxation rates, which show substantial differences and reduced *χ*^2^ values that are about two orders of magnitude greater than in the situation described above (177, 156 and 135 for ***R***(***D**_z_*), ***R***(***D**_y_*) and 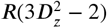, respectively) (Fig. 4A-C). We can improve the agreement by applying the reweighting procedure, but in contrast to the situation described above, it is more difficult to obtain a good agreement. Thus, when selecting *θ* to give *ϕ_eff_* = 0.2 we reduce the *χ^2^* values by about two-fold, compared to the about 10-fold in the case described above. We observe a similar behaviour also when only subsets of rate types are employed for fitting (Fig. S4). Nevertheless, it is clear that by changing the weights of the 1500 blocks we can improve the agreement between ‘experiment’ and simulations (Fig. 4A-C). Looking at the different methyl groups, we do not find that any specific residue gives rise to substantially worse agreement than others.

**Figure 4.**
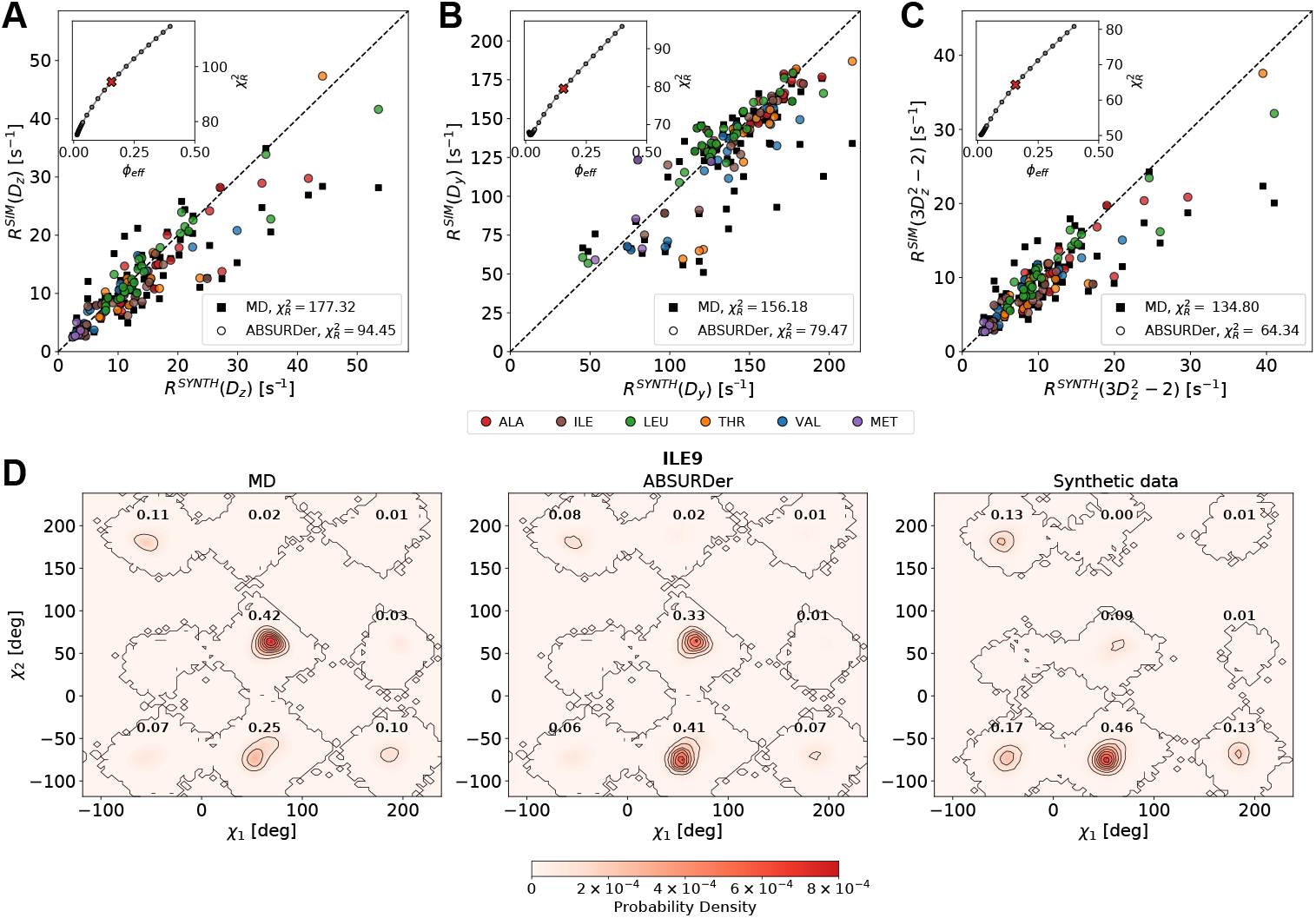
ABSURDer applied to synthetic data generated with different force fields. (A - C) ***R***(***D**_z_*), ***R***(***D**_y_*) and 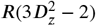 from both data sets are compared before (black squares) and after (filled circles) reweighting with ABSURDer. The rates are coloured by residue type. The insets show the behaviour of the respective reduced *χ*^2^ vs. *ϕ_eff_* during the reweighting using different values of *θ*. The chosen *θ* = 4500 corresponding to *ϕ_eff_* ≈ 0.2 is shown as a red cross. (D) Probability density distribution of the *χ*_1_ and *χ*_2_ angles of ILE9 before and after applying ABSURDer, and the corresponding ‘experimental’ data. Numbers in bold indicate probabilities of the corresponding rotamer states.

We again examined the consequences of the reweighting on the dihedral angles distributions, in this case focusing on ILE9 (Fig. 4D). The NMR relaxation experiments probe both the dynamics of the C^*γ*^ and C^*δ*^ in isoleucine residues, and the *χ*_1_ and *χ*_2_ dihedrals may display substantial couplings. We thus calculated the two-dimensional probability densities before and after reweighting and compared to the distribution from the Amber ff15ipq simulations that we used to generate the synthetic data. Although systematic differences remain between the reweighted and ‘experimental’ probability densities, it is clear that ABSURDer substantially corrects the probability densities. For example, the dominant rotamer in the Amber ff15ipq simulation used to calculate the synthetic NMR relaxation has *χ*_1_ as gauche^+^and *χ*_2_ as gauche^−^ and has a population of 0.46. The Amber ff99SB*-ILDN simulation has a population of 0.25 for this rotamer, a value that increases to 0.41 after reweighting. At the same time, the population of the gauche^+^/gauche^+^rotamer, which is almost not populated in the Amber ff15ipq simulation, decreases from 0.42 to 0.33. We note that although it is not unexpected, it is not trivial that a reweighting procedure employing kinetic data is able to correct static quantities such as equilibrium probability density. As above, we also find improvements in the backbone dihedral angles (Fig. S6). This example shows how the application of ABSURDer may help correct for inaccurate rotamer distributions in the simulations, though how much depends both on the information in the experimental data, and the sampling of the prior (force field). Indeed, no population shift can be obtained through reweighting if one of the rotameric states of interest is never observed during a simulation.

### Fitting experimental data globally

Having used synthetic data to show how ABSURDer makes it possible to fit MD simulations using NMR relaxation experiments, we now proceed to fit to experimental data on T4L. Before discussing the results, however, some comments are in order. In both cases above where we employed synthetic data to mimic experimental NMR relaxation rates, we used relaxation rates for all 100 methyl groups in T4L (apart from ALA146). In practice, however, not all of these can be measured accurately e.g. due to overlapping peaks or artifacts from strong ^13^C-^13^C coupling in certain residues (***Hoffmann et al., 2018b***). Thus, below we use only the 73 methyl groups whose relaxation rates were recently measured (***Hoffmann et al., 2018b***). To examine the effect of optimizing against a subset of residues we first repeated the reweighting procedure using the synthetic data, but restricting to the set of 73 methyl groups, and used the remaining 27 as a test set (Fig. S7). For both sources of synthetic data we find that optimizing on the 73 methyl groups lead to improvements in the remaining 27, unless aggressive reweighting to low *ϕ*_eff_ is employed.

We thus proceeded to use the previously measured data (***Hoffmann et al., 2018b***) recorded at 950 MHz, and compared these to the NMR relaxation parameters from the three 5 μs simulations, calculated as averages over the 1500 10 ns blocks (Fig. 5A–C). We find a reasonable agreement (reduced *χ*^2^ values of about 119, 179 and 92 for ***R***(***D**_z_*), ***R***(***D**_y_*) and 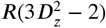, respectively), in line with similar calculations previously reported from ten 0.3 μs simulations. In contrast to the results described above for the synthetic data we observe, however, some systematic differences with the calculated rates on average being 16% lower than experiments.

**Figure 5.**
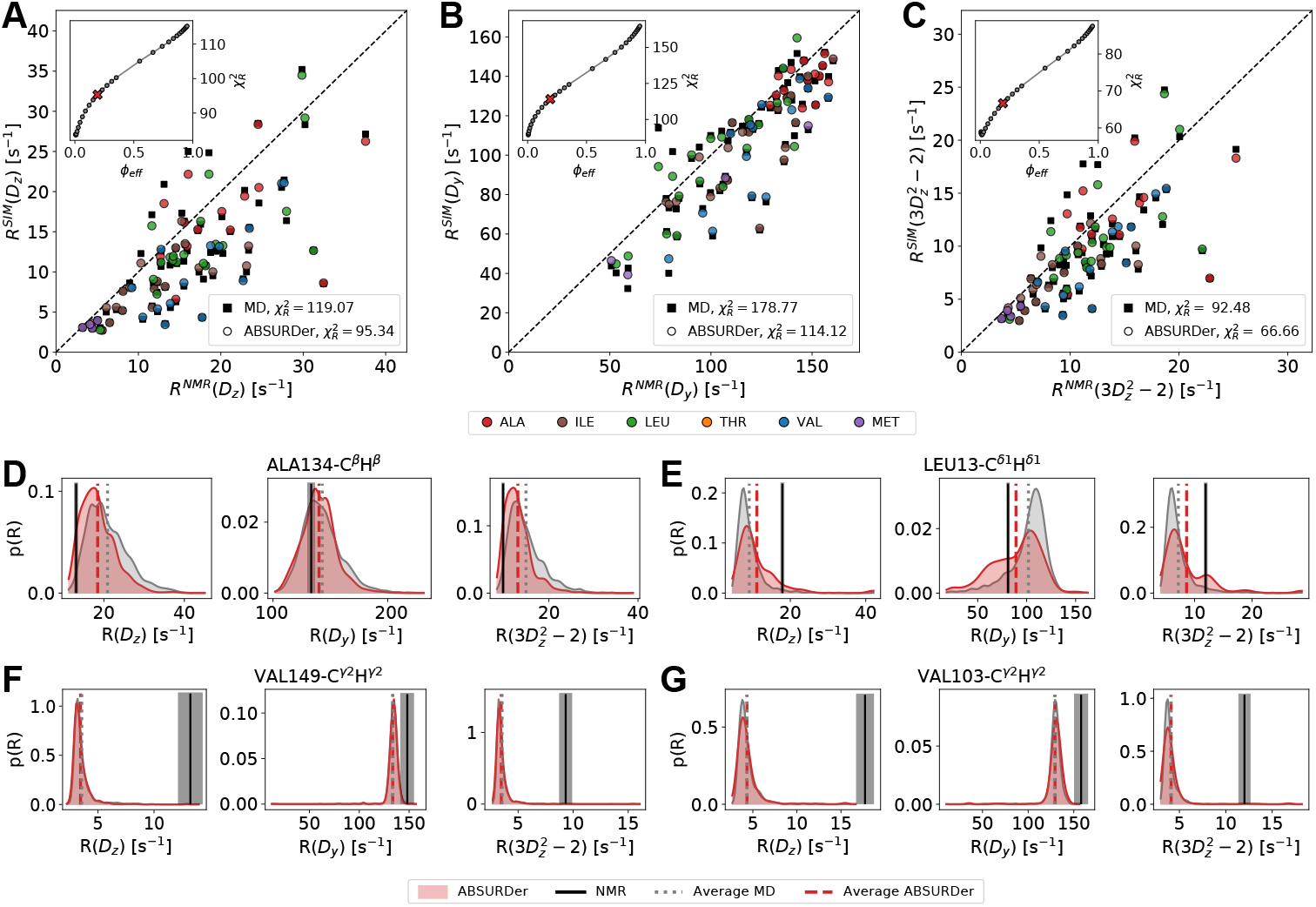
ABSURDer applied to experimental data. (A - C) ***R***(***D**_z_*), ***R***(***D**_y_*) and 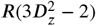 from both data sets are compared before (black squares) and after (filled circles) reweighting with ABSURDer. The rates are coloured by residue type. The insets show the behaviour of the respective reduced *χ*^2^ vs. *ϕ_eff_* during the reweighting using different values of *θ*. The chosen *θ* = 1400 corresponding to *ϕ_eff_* ≈ 0.2 is shown as a red cross. (D-G) Distributions of ***R***(***D**_z_*), ***R***(***D**_y_*) and 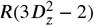 over the blocks for ALA134-C^*β*^H^*β*^ (D), LEU13-C^*δ*1^H^*δ*1^(E), VAL149-C^*γ*2^H^*γ*2^(F) and VAL103-C^*γ*2^H^*γ*2^(G) before (grey) and after (red) reweighting with ABSURDer. Vertical lines represent the average relaxation rate for the respective data set (NMR: solid black; MD: dotted grey; MD after ABSURDer: dashed red

We applied ABSURDer to improve agreement with the NMR experiments. When fitting to *ϕ_eff_* = 0.2 we are only able to obtain modest improvements of the calculated rates, decreasing the *χ*^2^ values by on average 30%. The difficulty in fitting this dataset also occurs when fitting only a single or two of the three types of relaxation rates, and as above ***R***(***D**_y_*) appears to be less correlated with the other two rates (Fig. S8). We also examined the agreement for each type of amino acid and find that in particular relaxation rates in valine and methionine methyl groups are poorly estimated and difficult to reweight (Fig. S9).

We then asked the question why some but not other relaxation rates and methyls can be improved by ABSURDer. Reweighting approaches such as ABSURDer rely on finding a sub-ensemble (or subset of trajectories) that fits the experimental data better than the full ensemble when averaged uniformly. Thus, the successful application requires that the calculated rates for the different blocks can be combined (linearly) to fit the data. We thus calculated the distribution of the relaxation rates over the 1500 blocks for selected methyl groups and compared the averages before and after reweighting to the experiments (Fig. 5D-G). For some residues and relaxation rates, exemplified by ALA134-C^*β*^H^*β*^ and LEU13-C^*δ*1^H^*δ*1^ in Fig. 5D-E, we find a relatively broad distribution where the relaxation rates calculated for the different blocks overlap with the experimental rates. In these cases it is possible to improve agreement by increasing the weights of some blocks and decreasing the weights of others. In several other cases, exemplified by VAL149-C^*γ*2^H^*γ*2^ and VAL103-C^*γ*2^H^*γ*2^ in Fig. 5F-G, the experimental relaxation rates lie outside the range sampled in our (blocks of) MD simulations. In such cases, reweighting cannot improve agreement substantially, because no linear combination of the blocks can get close to the experiments.

### Fitting subsets of experimental data

The problem of insufficient overlap is amplified when multiple rates are fitted simultaneously. If the dynamics at one site is uncorrelated with that at another site, there is no reason to expect that sub-trajectories that fit well the data for one methyl will do so for another. We note here that methods such as ABSURDer are fundamentally different from the standard approaches for analysing relaxation data. In most approaches to analyse NMR relaxation data, the relaxation parameters are fitted individually for each methyl, using only a global model for the rotational motion of the protein. In contrast, ABSURDer aims to find a set of trajectories that, at the same time, describe all relaxation rates at all sites. While this approach has the potential to quantify coupling between motions at different sites, the global analysis also potentially suffers from the ‘curse of dimensionality’.

In an attempt to alleviate the problems observed when fitting all relaxation parameters simultaneously (Fig. 5), we thus fitted various subsets of methyls. In particular, we (i) separately reweighted each type of methyl group and (ii) systematically tried to fit only those methyls for which there is some overlap between the distribution of calculated rates and experiments (as illustrated in Fig. 5D-G).

We thus fitted each type of methyl (either as a class of within each type of amino acid) individually. (Fig. 6A). For each class, we compare the agreement with experiment 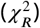 either before reweighting, when reweighting against the full set of methyls and when fitting only the specific set of methyls. As expected, it is possible to improve the fit substantially more when fitting just a single subset of methyls. We find that some types of methyls can be improved more than others, and note that this may be caused e.g. by differences in accuracy of the force field, differences in magnitude and time scales for different methyls, as well as differences in the number of the various methyls. We find that fitting one subset of methyls does not lead to improvement in the agreement with the remaining methyls that were not fitted (Fig. S10). This is in line with the difficulty in the global fitting, and with the observation that there can be substantial heterogeneity in the dynamics even among neighboring residues (***Best et al., 2005***; ***Igumenova et al., 2006***).

**Figure 6.**
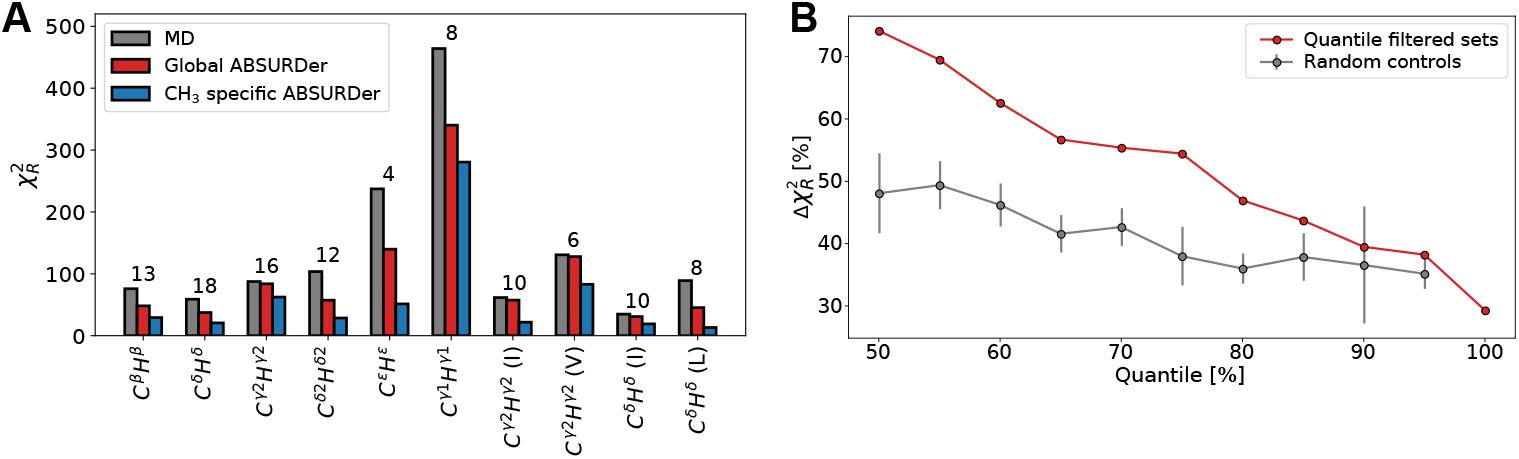
Fitting subsets of data using ABSURDer. (A) 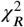 values obtained when comparing relaxation rates calculated from the MD simulations (grey) before and (blue) after reweighting to the experimental data for this specific class of methyls. For reference, we show in red the 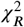 values obtained when all types of methyls are fitted as described in Fig. 5. Types of methyl groups that are present in more than one residue type were also analysed separately for each type of residue (I, V and L). The numbers over the bars indicate the number of methyl groups in that subset. (B) Relative change in the 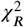 before and after reweighting as a function of the quantile threshold used to determine outliers (red). The grey points shows comparable results when we selected five subsets with the same number of methyls as in the corresponding red point. The error bars represent the standard error of the mean.

In addition to the difficulties that arise when fitting a large number of methyls simultaneously, reweighting methods such as ABSURDer are also based on the premise that accurate solutions are found within the set of sampled trajectories. We observed that the distribution of relaxation rates calculated over the 1500 different trajectories were unimodal (such as the examples in Fig. 5D–G)), but that the different methyls and rates differed on the amount of ‘overlap’ between the distribution and the average experimental rate. We thus asked the question whether it would be easier to fit rates with more such overlap and proceeded as follows: According to a fixed quantile threshold, we removed all those methyls for which at least one of the three experimental rates did not fall within the given quantile. This enabled us to systematically remove outliers by lowering the quantile threshold, and we thereby used ABSURDer to fit groups of methyls with increasing numbers of such outliers (Fig. 6B). As expected, the results show that it is possible to fit better (as quantified by the drop in 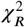 upon fitting) as the least overlapping methyls are removed. To control for the effect of fitting fewer methyls we applied ABSURDer to an equal-sized subset chosen randomly, and while smaller subsets enable better fitting the decrease in 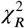 is substantially smaller than when outliers are systematically removed.

Together, the results obtained by applying ABSURDer on subsets of methyls provide strong additional evidence for the potential of the method and highlights future areas for improvement. The results show that, in principle, ABSURDer could be applied to reweight methyl subsets used to identify local properties of the protein, e.g. when one is interested in the dynamics of specific protein sub-regions or domains. Further, fitting will be aided by more extensive sampling (providing more samples in the tails of the distribution of rates) and improved force fields (moving the average of the distribution of rates closer to experiments).

## Conclusions

We have here presented ABSURDer, an extension of the ABSURD approach (***Salvi et al., 2016***), and applied it to NMR relaxation measurements of side chain dynamics. We have described and validated ABSURDer using synthetic data and applied it to experimental NMR data. As expected, for the three levels of ‘complexity’ we obtain different levels of agreement after reweighting between simulated and ‘experimental’ data.

When we used the same force field to generate synthetic data as that used to fit the data (ff99SB*-ILDN) we find good agreement even before reweighting, in line with the fact that the calculations of the relaxation parameters are relatively precise even using only a few microseconds of simulations (Fig. 2). Nevertheless, we can improve agreement further by reweighting and find, for example, improvements in dihedral angle distributions when we fit against the relaxation rates. As expected from the fact that we fit against the relaxation rates, we also find overall improvement of the spectral density functions after reweighting.

The overall observations were similar when we generated synthetic data using Amber ff15ipq and fitted using simulations generated with ff99SB*-ILDN. Here, we are still able to achieve a substantial improvement in the quality of the fit (around a 50% decrease in the overall *χ*^2^). Notably, also in this case the overall agreement between ‘experimental’ and simulated spectral densities and rotamer distributions increases upon reweighting.

When we fitted the experimental NMR data to the simulations with ff99SB*-ILDN we obtain a more modest improvement (an overall 30% decrease in *χ*^2^). Examining the probability distribution of rates over the different blocks and residues, we find that while some of these are broad and highly overlapping with the experimental average, many others are sharp and incompatible with the experimentally determined rates. Clearly, it is difficult to reweight when there is only little overlap between the experimental value and the rates calculated from the different blocks. This problem is exacerbated by the fact that we fit all rates simultaneously leading to a problem akin to the ‘curse of dimensionality’.

The natural question arises whether it is possible to increase the agreement between simulated and experimental NMR relaxation rates further. As described in the Introduction, the main sources of error between simulated and experimental data are: (i) insufficient sampling, (ii) force field inaccuracies, and (iii) remaining errors in the forward model used to estimate the relaxation rates from the trajectories. In the absence of a measure of ground truth it may, however, be difficult to disentangle these effects.

Our analysis of errors due to sampling (Fig. 2B-D) shows that the rates calculated from the 15 μs of simulation are rather precise compared e.g. to the deviation between experimental and calculated rates (Fig. 5). We have also calculated the total weight of the blocks coming from each of the three 5 μs simulations after reweighting, and while these sums differ from the expected average of 0.33, none of them are greater than 0.5 (Tab. S1). While these observations suggest that sampling is not the main source of deviation, the situation during reweighting is more complicated. Indeed, the selection of the block size demonstrates one of the complicating factors (Fig. 2A); using a too short block size leads to systematic errors while using too long blocks leads to ‘local convergence’ and thus a narrow distribution of rates across the blocks. While a 10 ns block size strikes a reasonable balance between these opposing factors it is clear that, in many cases, there is not sufficient overlap between the distribution of calculated rates and the experimental values (Fig. 5). Clearly, additional sampling to obtain more blocks could help alleviate this situation by increasing the number of samples in the tails of these distributions, though in many cases the deviations are so large that substantially more sampling would be needed. However, regardless of the amount of sampling, the choice to cut the trajectory into blocks naturally introduces a pre-averaging of the dynamics at the sub-nanosecond scale; therefore, if the dynamics at short timescales is inaccurate because of force field imprecision, there is nothing ABSURDer can do to fix it. One future solution to this problem could be to use blocks of different sizes and combine the derived data. In this way one could fit motions on fast timescales by using small blocks and slower motions with longer blocks. Such an approach could be used together with relaxation data from a wider range of magnetic field strengths to probe motions on a range of time scales (***Cousin et al., 2018***).

Our results on fitting subsets of experimental data demonstrate that one of the problems in the practical application of ABSURDer is the difficulty of constructing a single global model of the protein dynamics. This likely arises because many side chain motions are only weakly and locally coupled (***Best et al., 2005***) and that any potential long-range coupling could arise from these local effects (***Fenwick et al., 2011***; ***DuBay et al., 2011***; ***Lindorff-Larsen et al., 2016***). In addition to using simulation-derived correlation functions, the ABSURDer approach differs from standard methods for analysing NMR relaxation data by aiming to fit the dynamics of all residues at the same time. One potential way forward might be to create (structurally) local models of the dynamics (to overcome the high-dimensionality of the problem) and then reconstruct models of the full protein dynamics from these (***Olsson and Noé, 2019***; ***Hempel et al., 2021***).

As reweighting methods such as ABSURDer rely on an overlap between experiment and calculated parameters prior to reweighting, they are aided by good agreement before reweighting (***Hummer and Köfinger, 2015***; ***Orioli et al., 2020***; ***Larsen et al., 2020***). Indeed the extent of reweighting needed to obtain good agreement is related to a measure of the error in the force field (***Qian, 2001***; ***Orioli et al., 2020***). Thus, as starting point for our calculations we used force fields that have recently been shown to give improved agreement to side chain relaxation data (***Hoffmann et al., 2018a**,b, **2020***). Those studies used quantum-level calculations to modify the barriers for the methyl spinning, and demonstrated improved agreement with experimental NMR relaxation rates. More specifically, it was shown that methyl spinning barriers where generally too high, and that a small decrease (obtained by fitting to coupled cluster quantum chemical calculations of isolated dipeptides) lead to increased spinning rates and better agreement with experimental data in both T4L and ubiquitin (***Hoffmann et al., 2018a**, b, **2020***). Other effects might, however, contribute to the imperfect agreement between experiments and simulations. For example, the quantum calculations did not include information about how methyl spinning rates are influenced e.g. by side-chain packing or tunneling effects (***Chatfield and Wong, 2000***; ***Chatfield et al., 2004***).

A final source of error between experiment and simulation that may also prevent reweighting is the accuracy of the forward model used to calculate the relaxation rates from simulations. Here we have used a recently described approach to calculate deuterium relaxation rates from MD simulations (***Hoffmann et al., 2018a**, b*). While the approach is highly accurate, there are some sources of errors including for example the treatment of tumbling motions, non-symmetry of partially deuterated methyl groups, finite sampling of the fastest dynamics (we save frames every 1 ps) and fitting of the correlation functions to a fixed number of exponentials. We note here that we simulate a fully protonated molecule, and while there may be effects of (partial) deuteration on the local structure and dynamics these are expected to be small (***Lee et al., 1999***; ***Ishima et al., 2001***; ***Mittermaier and Kay, 2002***). One potentially relevant effect is the difference in viscosity between the (H_2_O) water model used in simulations and the mixture of H_2_O and D_2_O that is used in NMR experiments (***Hoffmann et al., 2018a***), though this mostly would have an effect when modelling the rotational motions from simulation.

In summary, we have presented ABSURDer and applied it to study motions in methyl containing residues in a folded protein. The approach is general and may be extended to other types of residues and nuclei, to other NMR parameters that depend on relaxation rates such as nuclear Overhauser enhancements, and even to data beyond those generated by NMR such as e.g. fluorescence correlation and neutron spin-echo spectroscopy.

## Methods

### Sampling

The starting structure for our MD simulations was the X-ray structure of the cystein-free T4L SER44GLY mutant (PDB 107L) (***Blaber et al., 1993***) where we changed GLY44 back to a serine. We performed three different sets of simulations: Three 5 μs long (“MD Data” in all three analyses) and five 1 μs long (“NMR Data” in the first synthetic data set) simulations, both using the AMBER ff99SB*-ILDN / TIP4P-2005 protein force field (***Lindorff-Larsen et al., 2010***; ***Hornak et al., 2006***; ***Best and Hummer, 2009***), and three 1 μs long simulations using the AMBER ff15ipq / SPC/E_*b*_ (***Debiec et al., 2016***) force field (“NMR Data” in the second synthetic data set). We applied a modification to the methyl rotation barriers to both force fields as recently described (***Hoffmann et al., 2018a**, **2020***). In the case of the AMBER ff15ipq simulation, we used the Amber input preparation module LEAP from AmberTools17 (***Case et al., 2017***) to set up the system and we afterwards converted the topology to GROMACS file format using ParmED (***Swails et al., 2010***).

All MD simulations were carried out with GROMACS v2018.1. We used a periodic truncated dodecahedron of 400 nm^3^ volume as a simulation box, keeping 1.2 nm distance from the protein which was centered in the cell. We solvated the system with 12230 TIP4P-2005 (***Abascal and Vega, 2005***) water molecules and 12247 SPC/E_*b*_ water molecules, respectively, retaining crystal waters. To neutralize the system and simulate it in a physiological salt concentration, we also added 150 mmol L^−1^Na^+^/Cl^−^ ions. We minimized the systems with 50000 steps of steepest descent and equilibrated them for 200 ps in the *NPT* ensemble, using harmonic position restraints (force constants of 1000 kJ mol^−1^ nm^−2^) on the heavy atoms of the protein. Equations of motion were integrated using the leap-frog algorithm. We set a cut-off of 1 nm for the Van der Waals and Coulomb interactions and employed Particle-Mesh Ewald summation for long-range electrostatics (***Essmann et al., 1995***) with 4^*th*^ order cubic interpolation and 0.16 nm Fourier grid spacing. We ran the production simulations in the *NPT* ensemble, using the velocity re-scaling thermostat (***Bussi et al., 2007***) with a reference temperature of 300 K and thermostat timescale of *τ*_T_ = 1 ps, and the Parrinello-Rahman barostat with a reference pressure of 1 bar, a barostat timescale *τ*_P_ = 2ps and an isothermal compressibility of 4.5 × 10^−5^bar^−1^. Finally, we saved the protein coordinates every 1 ps and, after running the simulations, we removed the overall tumbling of the protein by fitting to a reference structure.

### Spectral Density Mapping

We used a previously described approach (***Hoffmann et al., 2018b***) to calculate NMR relaxation rates from computed spectral densities. The original work used multiple 300 ns-long simulations, however, we analysed our simulations in a block-wise fashion, using 10 ns long, non-overlapping blocks. This yielded 1500 blocks for the set of 3x 5 μs-long simulations, 500 blocks for the set of 5x 1 μs-long simulations and 300 blocks for the 3x 1 μs long simulations. We started by calculating the internal time correlation function (TCF), *i.e*. without global tumbling, of the three 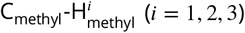 bond vectors up to a maximum lag-time of 5 ns for all side-chain methyl groups. Next, we fitted the internal TCFs with six exponential functions and an offset,

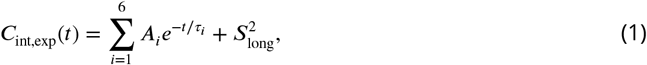

where 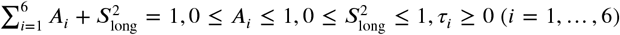 and 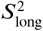 is the long-time limit order parameter. We note that one should not attribute physical meaning to these amplitudes and timescales (***Bremi et al., 1997***). Since the protein was found to be best represented by an axially symmetric tumbling model (***Hoffmann et al., 2018b***), we introduced methyl-specific global tumbling times *τ_R,i_* by multiplying the internal TCF with a single-exponential thus, yielding the total TCF:

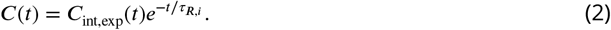

In this case, we applied the experimental methyl group-specific tumbling times, which, for the synthetic data sets, we extracted based on analysing backbone N-H dynamics (***Maragakis et al., 2008***; ***Hoffmann et al., 2018b***). Briefly, we calculated TCFs for the backbone N-H bond vectors and fitted them to the three-parameter Liparo-Szabo model:

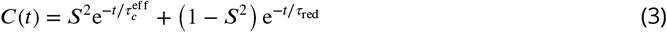

with 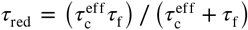 and where *S*^2^, *τ_f_* and 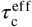 are used as fitting parameters. Next, we used the fitted rotational tumbling times and the initial structure of the protein, translated so that its center of mass is located at the origin (PDBinertia, http://comdnmr.nysbc.org), to calculate the principal axis frame (Quadric (***Lee et al., 1997**)*). Finally, we extracted the principal values ***D**_xx_, **D**_yy_* and ***D**_zz_* from the diffusion tensor and calculated the methyl-specific rotational diffusion constants 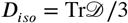. Following that, we projected the C-C_methyl_ vectors onto the principal axes of ***D***, ***D_i_*** = ***D***_iso_ − ***P***_2_(cos *θ*)(***D**_zz_* − ***D**_yy_*)/3 (according to an axially symmetric diffusion model) and we used ***D_i_*** to calculate the methyl-specific tumbling times *τ_R,i_* = 1/(6***D**_i_*).

After introducing *τ_R,i_* we transformed the total TCF into a spectral density function:

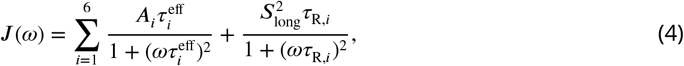

where 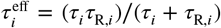. Finally, we calculated the relaxation rates directly from ***J***(0), ***J***(*ω*_D_), ***J***(2*ω*_D_) via

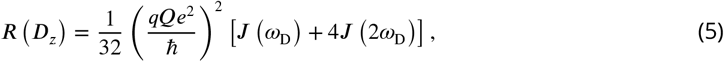

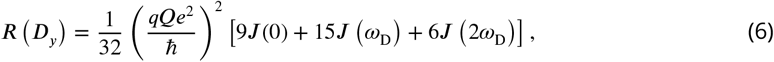

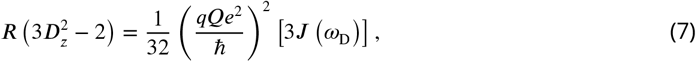

where 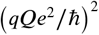 is the quadrupolar coupling constant of deuterium. We used 145.851 MHz as the Larmor frequency *ω*_D_ of deuterium for all data sets which corresponds to a Bruker magnetic field strength of 22.3160T that was also used to measure the experimental NMR relaxation rates.

We used this approach for each side-chain methyl group and each of the 1500 blocks to calculate sets of the three relaxation rates, which we employed as a input to the ABSURDer reweighting. When generating synthetic data, we calculated average rates over the blocks, and we used the standard error of the mean over the blocks as the errors when calculating *χ*^2^ and for reweighting with ABSURDer.

### Reweighting

Our reweighting approach builds upon the previously described ABSURD method (***Salvi et al., 2016***),where a longer set of trajectories are divided into *blocks*, each of which is long enough to estimate the relaxation rates of interest. Finally, a weighted average of the rates is performed over the blocks, where the weights, **w**, are obtained through the optimisation of a functional which determines the agreement between simulated and experimental rates. In particular, the ABSURD functional is given by

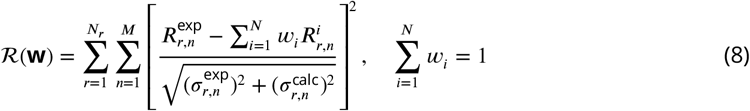

where *r* ranges over the *N_r_* types of experimental NMR relaxation rates, ***M*** is the overall number of measured rates and *N* represents the number of blocks which the trajectory has been cut into, 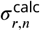 is the experimental error on the *n*-th rate of the *r*-th type, 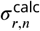 is the standard error on the simulated rates, estimated by averaging all the *n*-th rates of the *r*-th type over the different blocks. The optimal set of weights is obtained as the corresponding minimum in weight space, 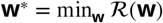. Each weight is associated to a block from the trajectory and it encodes the relevance of each block to the dynamical ensemble.

The functional in Eq. 8 is unregularised, which sometimes may lead to overfitting (***Orioli et al., 2020***). In the original implementation of ABSURD, this was avoided by extensive cross-validation using multiple relaxation rates, measured at multiple magnetic fields. A complementary approach is to use the MD simulations as a statistical prior and to balance the trust in the data and the experiments (***Orioli et al., 2020***).

We take this latter approach by introducing an extension to the ABSURD functional (which we denominate ABSURDer) to make it less prone to overfitting. In particular, we add a Shannon relative-entropy regularization ***S***(**w**) (***Kullback and Leibler, 1951***; ***Gull and Daniell, 1978***) and the new functional takes the following form:

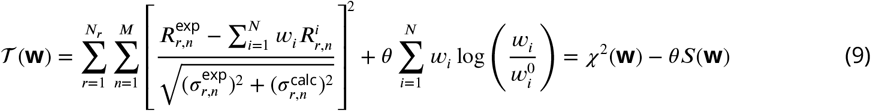

where 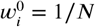 represent the initial, uniform weights provided by the prior. The parameter *θ*, is used to set the balance between our trust in the prior and in the experimental data. In particular, for *θ* → ∞, an infinite trust is put on the prior and the weights *w_i_* stay at their initial values. In the opposite limit, *θ* → 0, all the trust is put on the experimental data and the weights are allowed to change freely to accommodate this trust. In this case, the functional 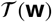 reduces to that used in ABSURD. The two limits of *θ* can be seen also from parameter *ϕ*_eff_(**w**) = exp(*S*(**w**)), which provides a measure of the effective fraction of blocks retained for a given set of weights. In particular, for *θ* → ∞we have *ϕ*_eff_(**w**) = 1, as 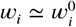. On the other hand, for *θ* → 0 and in the case where the prior is very different from the experimental data, *ϕ*_eff_(**w**) may take on very small values.

We minimised functional 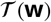 using the limited memory Broyden-Fletcher-Goldfarb-Shanno algorithm (L-BFGS-B) in its SciPy implementation (***Virtanen et al., 2020***). In each reweighting run we employed 30 values of *θ* selected in the range [0,8000]. To accelerate convergence of the optimization, we started by optimising the *θ* = 8000 functional using **w**_0_ as a guess set of weights, then we employed the obtained optimal weights as guess for the successive minimisation run and so on, iteratively. We stress that the value of **w**_0_ in Eq. 9 remained unchanged throughout the minimisation, regardless of the choice of the guess input weights. The ABSURDer code is available under a GNU GPL v3.0 license at github.com/KULL-Centre/ABSURDer and scripts and data to generate the results in this paper are available at github.com/KULL-Centre/papers/tree/master/2020/ABSURDer-kummerer-orioli-et-al-2020.

## Acknowledgments

We are grateful to Profs. Lars V. Schäfer and Frans A.A. Mulder for discussions, help and comments on the manuscript. We acknowledge support by a grant from the Lundbeck Foundation to the BRAINSTRUC structural biology initiative (155-2015-2666, to K.L.-L.), the NordForsk Nordic Neutron Science Programme (to K.L.-L.), the Carlsberg Foundation (CF17-0491, to Y.G.), and the Novo Nordisk Foundation (NNF15OC0016360, to K.T. and K.L.-L.). We acknowledge access to computational resources from the ROBUST Resource for Biomolecular Simulations (supported by the Novo Nordisk Foundation grant no. NF18OC0032608).

## Supporting Material

**Supporting Figure 1.**
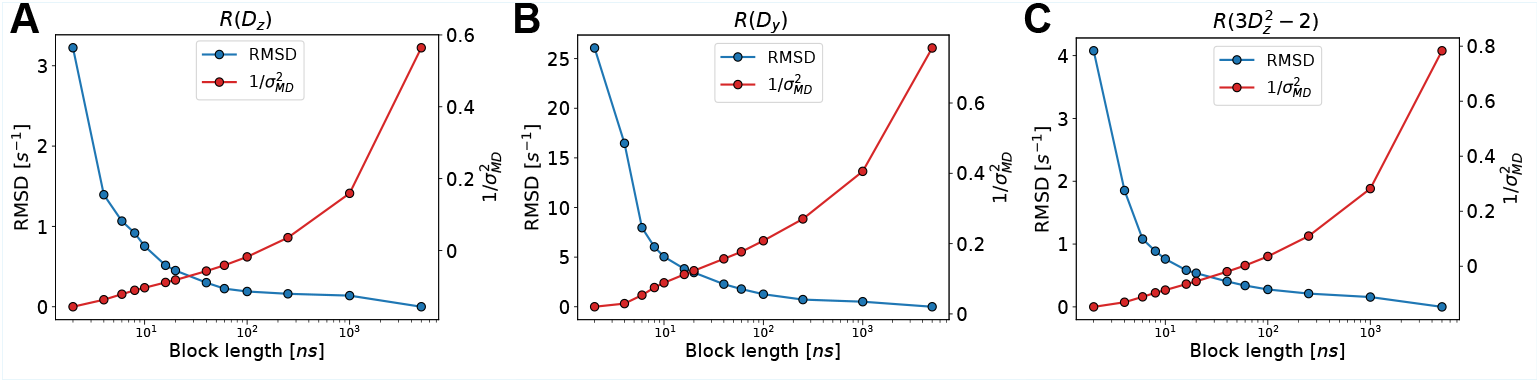
Assessment of convergence and effect of block size. RMSD of NMR relaxation rates (blue) and average inverse variance with respect to the full MD dataset (red) is shown as a function of the block length.

**Supporting Figure 2.**
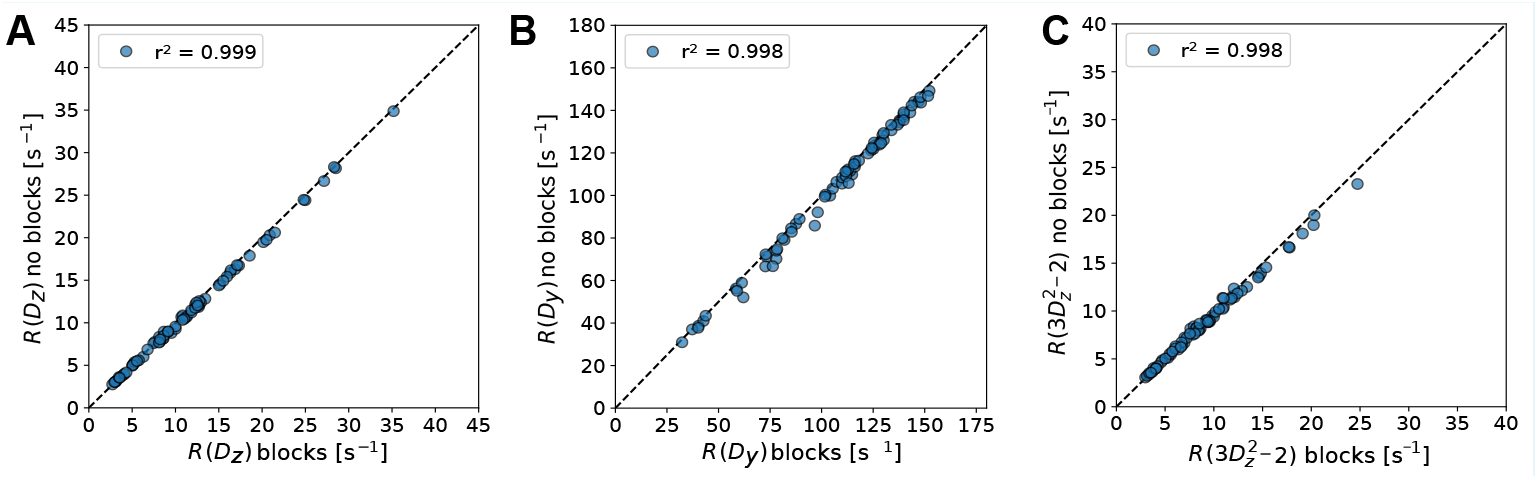
Comparison of NMR relaxation rates calculated from 1500 10ns blocks and the full MD dataset. (A) ***R***(***D**_z_*), (B) ***R***(***D**_y_*) and (C) 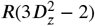 calculated from the MD dataset using 1500 10ns long blocks and three 5 μs long blocks (no blocks). The label shows the corresponding Pearson correlation coefficient.

**Supporting Figure 3.**
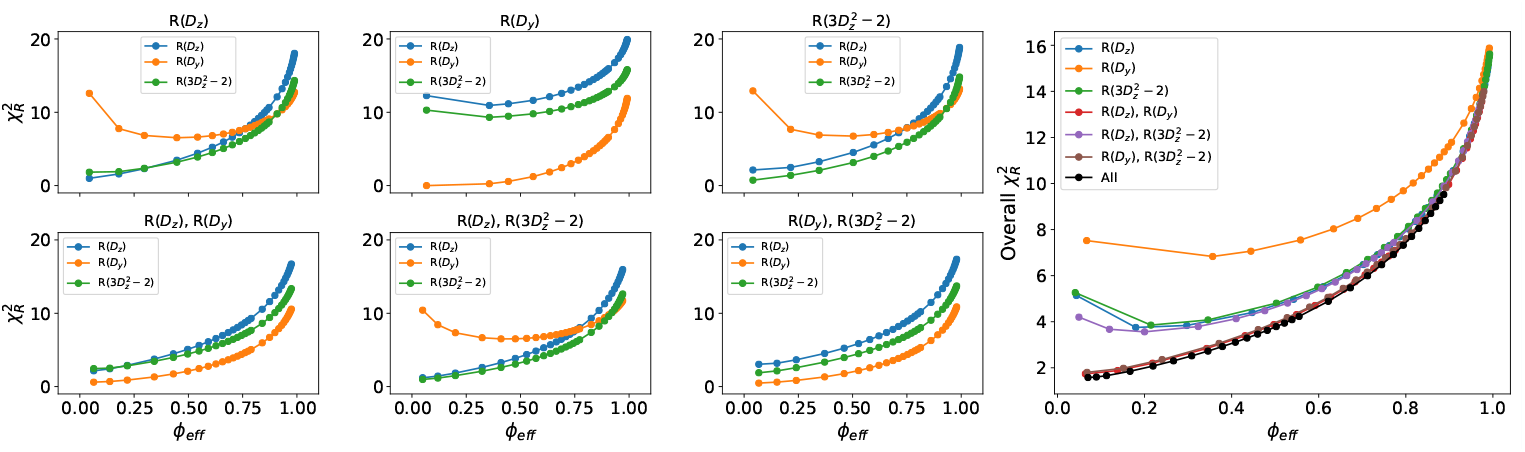
Cross validation of ABSURDer reweighting when synthetic data were generated using the Amber ff99SB*-ILDN force field. The panels show 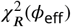 curves for each of the three types of NMR relaxation rates. Each panel differs by which data was used in reweighting (label above panel), and we show the results using all six possible combinations of the three rates with the large panel corresponding to all three rates.

**Supporting Figure 4.**
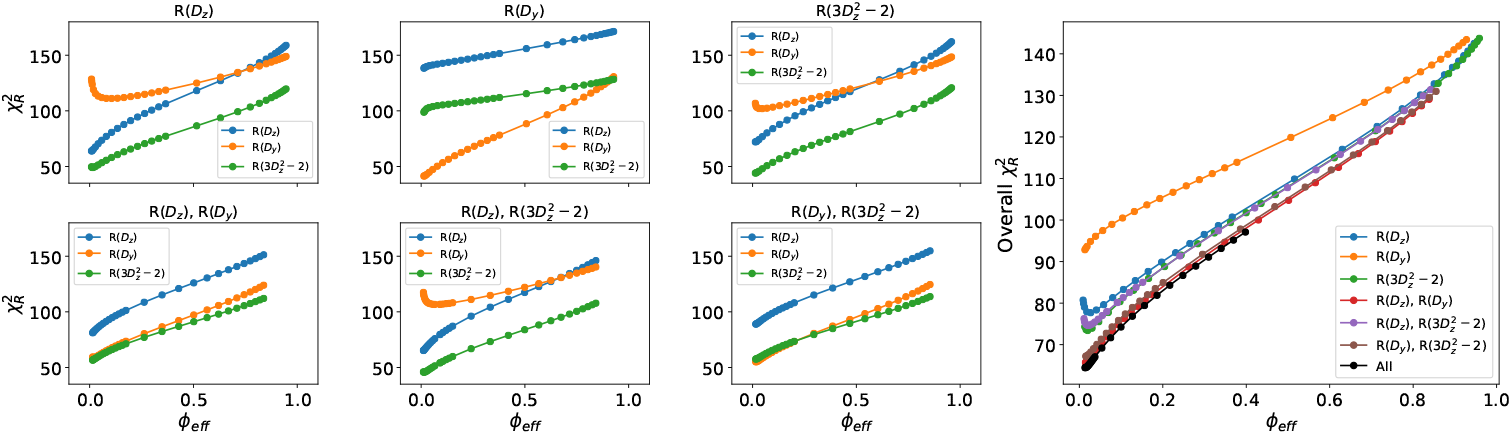
Cross validation of ABSURDer reweighting when synthetic data were generated using the Amber ff15ipq force field. The panels show 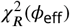 curves for each of the three types of NMR relaxation rates. Each panel differs by which data was used in reweighting (label above panel), and we show the results using all six possible combinations of the three rates with the large panel corresponding to all three rates.

**Supporting Figure 5.**
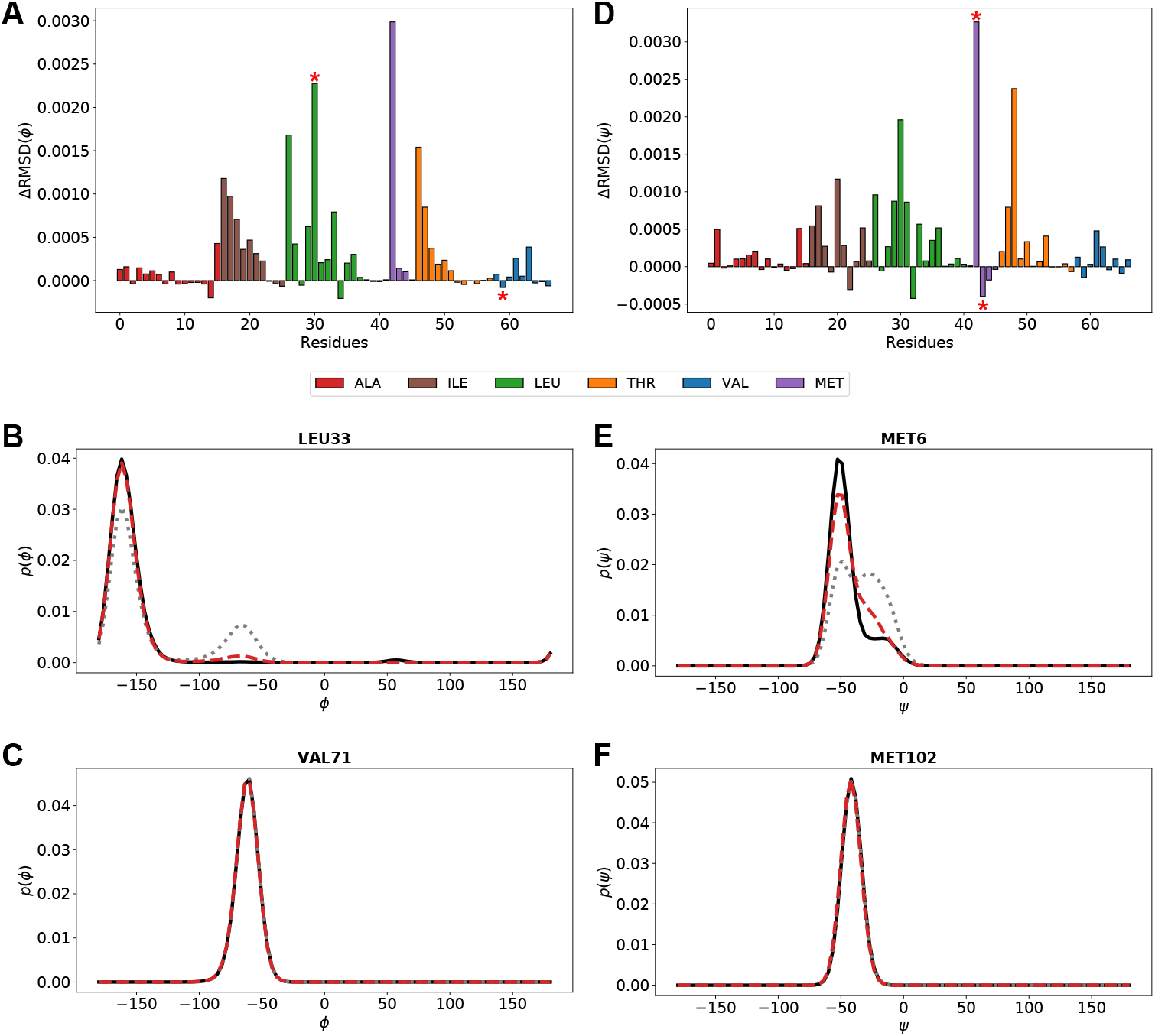
Effect of ABSURDer reweighting on backbone dihedral angle distributions. Difference RMSD (ΔRMSD) between distributions for the backbone (A) *ϕ* and (B) *ψ* dihedral angles. We calculated the RMSD of the dihedral angle distributions before (grey dotted line) and after (red dashed line) reweighting, in both cases using the distribution used to generate the synthetic data generated by Amber ff99SB*-ILDN as the reference (black solid line). Thus, large positive values of ΔRMSD indicate highly improved distributions after reweighting whereas negative values indicate distributions worsened by the application of ABSURDer. The bars are colored by residue type and asterisks mark residues highlighted in panels C-F. Examples of residues for which ABSURDer (C,D) substantially improve the dihedral angle distributions and (E,F) worsens them.

**Supporting Figure 6.**
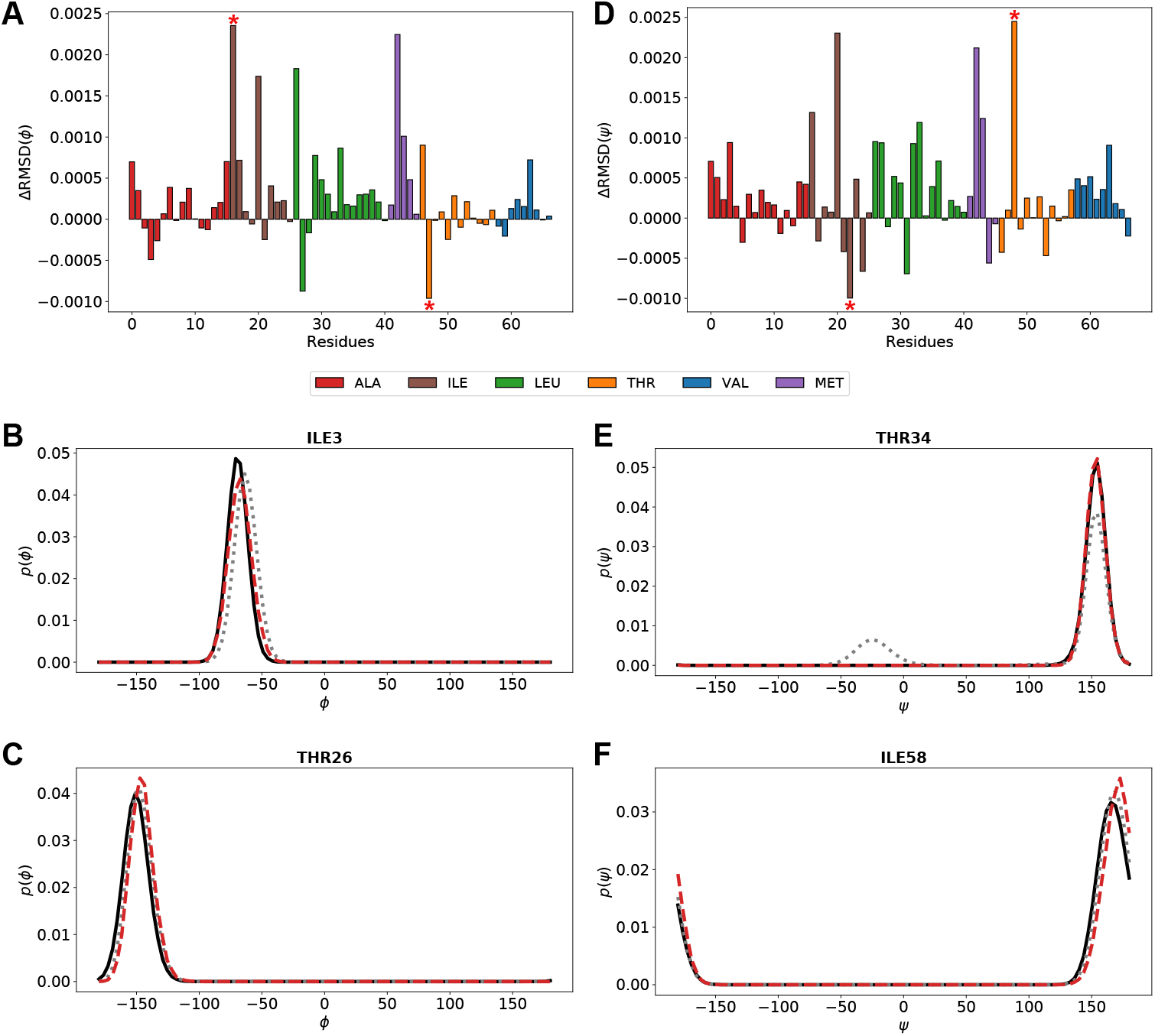
Effect of ABSURDer reweighting on backbone dihedral angle distributions. Difference RMSD (ΔRMSD) between distributions for the backbone (A) *ϕ* and (B) *ψ* dihedral angles. We calculated the RMSD of the dihedral angle distributions before (grey dotted line) and after (red dashed line) reweighting, in both cases using the distribution used to generate the synthetic data generated by Amber ff15ipq as the reference (black solid line). Thus, large positive values of ΔRMSD indicate highly improved distributions after reweighting whereas negative values indicate distributions worsened by the application of ABSURDer. The bars are colored by residue type and asterisks mark residues highlighted in panels C-F. Examples of residues for which ABSURDer (C,D) substantially improve the dihedral angle distributions and (E,F) worsens them.

**Supporting Figure 7.**
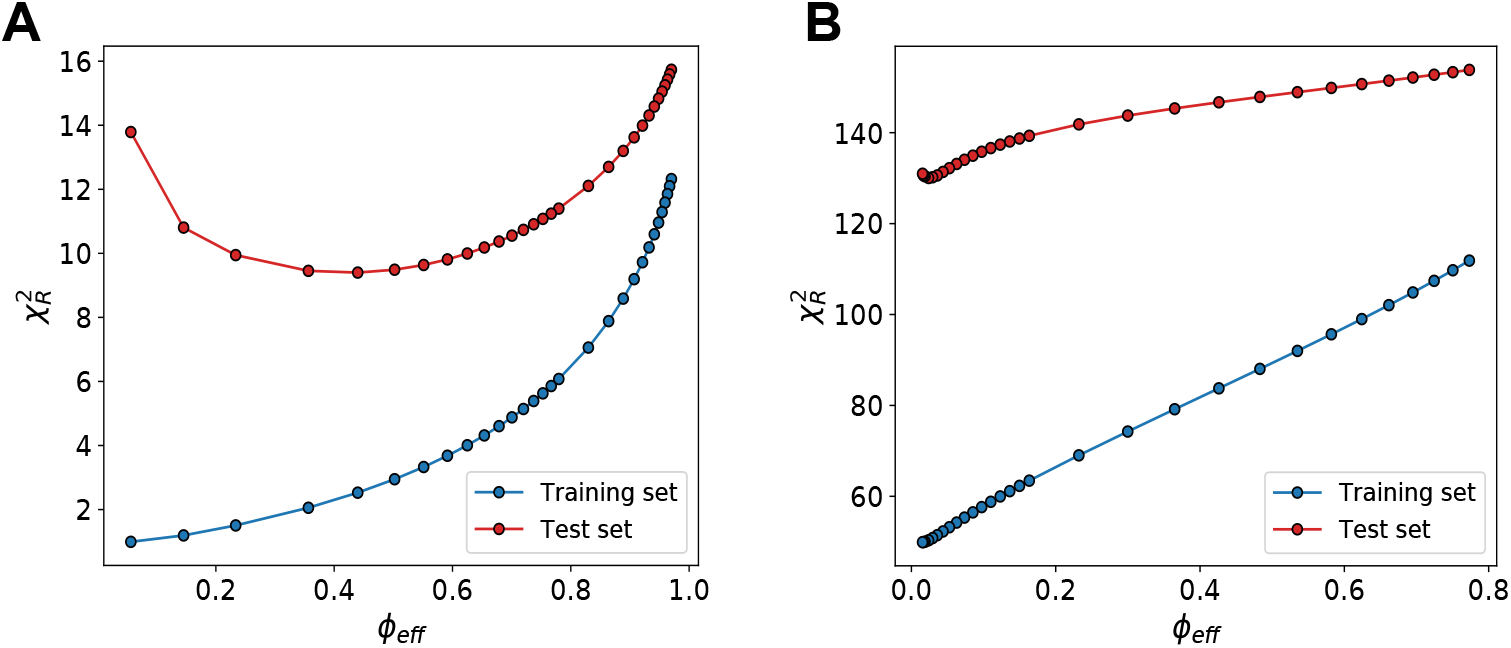
Effect of ABSURDer reweighting on NMR relaxation rates not included in the optimisation. We used ABSURDer with synthetic data to fit the relaxation rates for the (blue) 73 methyl groups that have been probed experimentally, and (red) cross-validate across the remaining 27 methyl groups. (A) Synthetic data generated with Amber ff99SB*-ILDN. (B) Synthetic data generated with Amber ff15ipq.

**Supporting Figure 8.**
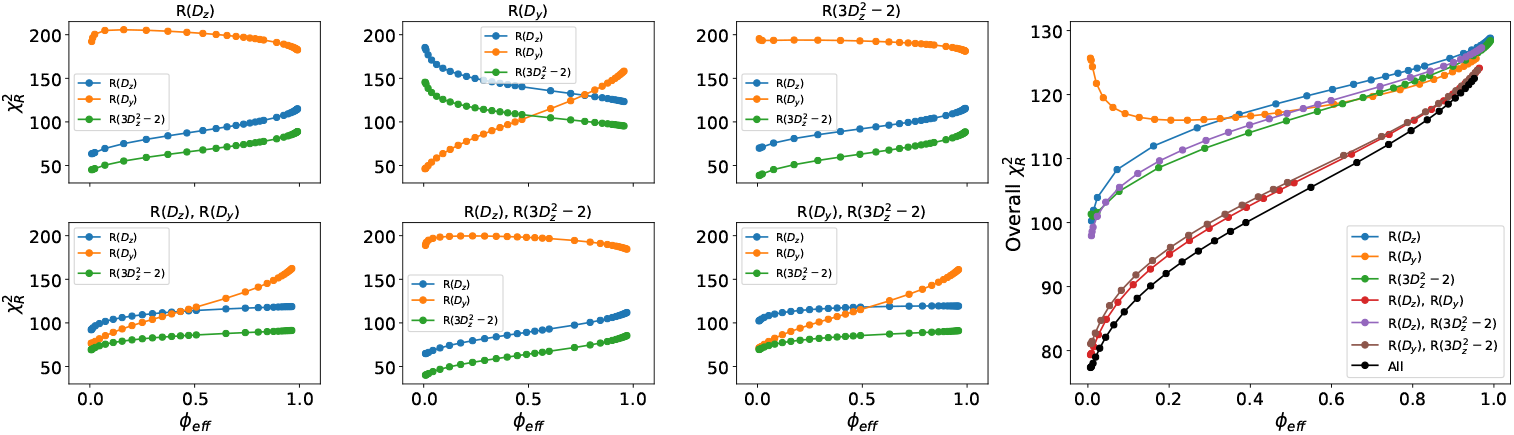
Effect of ABSURDer reweighting on experimental data when only a subset of NMR relaxation rate types are used. 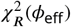 curves for the three kinds of rate separately obtained by fitting with respect to all the possible combinations of rate types (6 panels on the left) and the corresponding average (overall 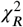, right panel).

**Supporting Figure 9.**
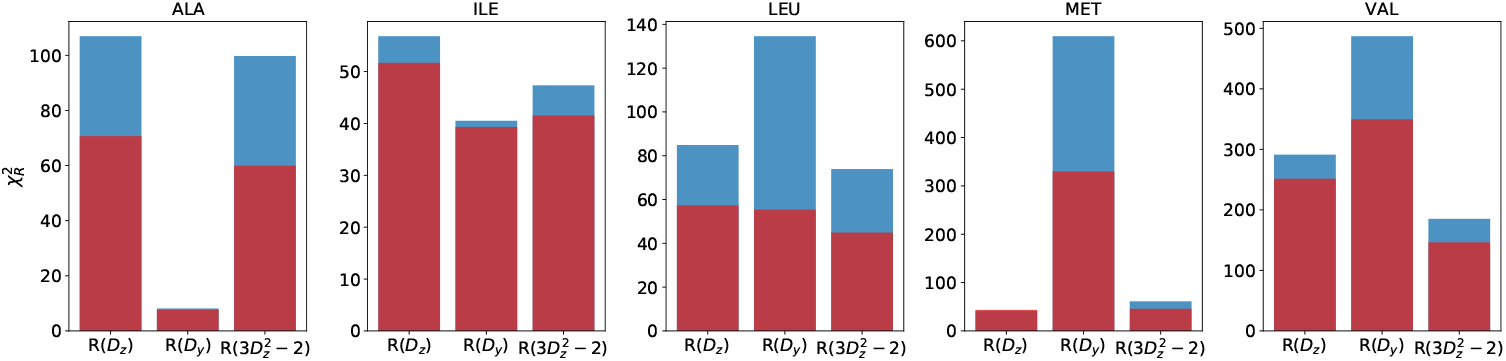
Effect of ABSURDer on the different residue types. Residue specific 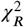 between calculated and experimental NMR relaxation rates before (blue) and after (red) reweighting. The results are obtained by averaging the residuals over the 15 ALA, 20 ILE, 5 MET, 14 VAL and 20 LEU residues. Note that each panel uses different scales for 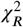.

**Supporting Figure 10.**
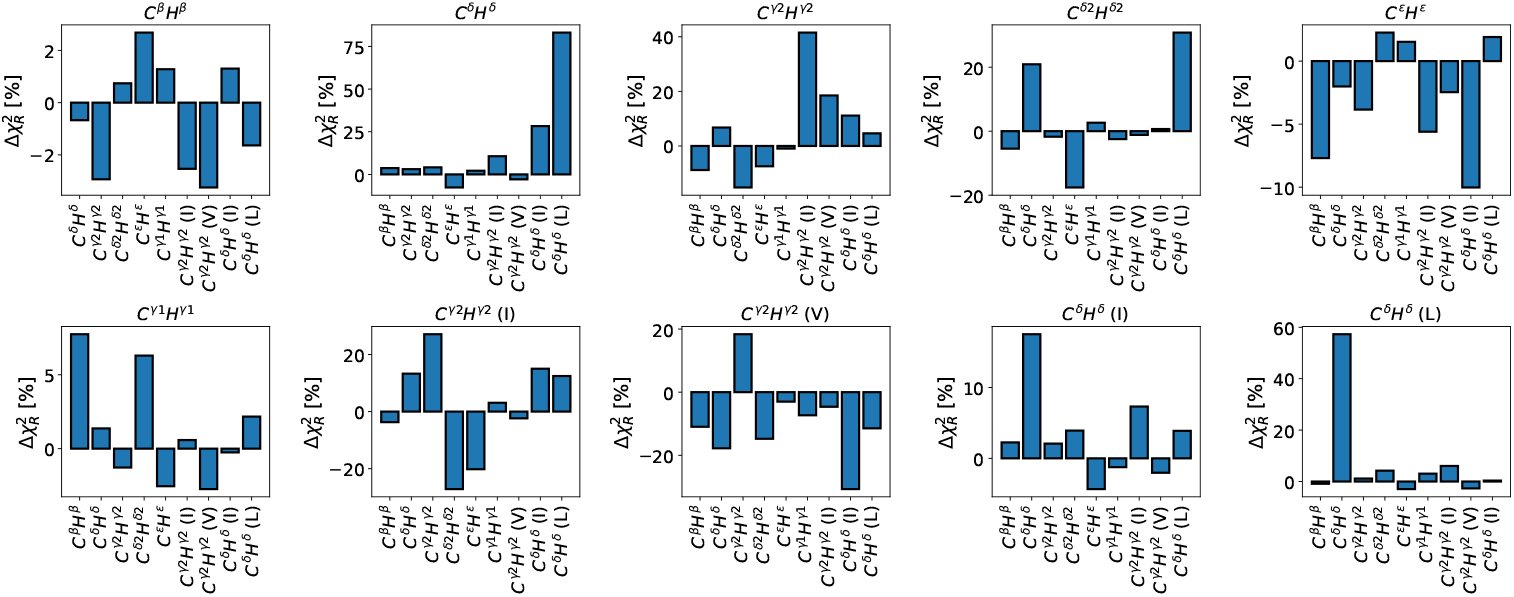
Result of ABSURDer reweighting on NMR relaxation rates that were not included in the optimisation of specific methyl subsets. We used ABSURDer to fit the relaxation rates for each methyl type separately and cross validated the results against the remaining methyl subsets. The y-axis reports the relative change in 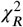 after the reweighting has been carried out on the methyl class specified in the plot titles. Negative values indicate that the agreement with experiments decreased, while positive values indicate improved agreement. Methyl types that are present across different types of residues are also shown separately (i.e. I, L and V).

**Supporting Table 1.** Sum of the weights for blocks in each of the three independent 5 μs long trajectories.

|  | Trajectory 1 | Trajectory 2 | Trajectory 3 |
| --- | --- | --- | --- |
| Same force fields | 0.21 | 0.5 | 0.29 |
| Different force fields | 0.28 | 0.45 | 0.27 |
| Experimental | 0.38 | 0.28 | 0.34 |

